# Sex differences in human musicality are negligible

**DOI:** 10.1101/2023.05.23.541970

**Authors:** Mila Bertolo, Daniel Müllensiefen, Isabelle Peretz, Sarah C. Woolley, Jon T. Sakata, Samuel A. Mehr

**Affiliations:** Integrated Program in Neuroscience, McGill University, Quebec, QC, Canada H1A 2B4; Center for Research on Brain, Language, and Music, Quebec, QC, Canada H3G 2A8; International Laboratory for Brain, Music, and Sound Research (BRAMS), Department of Psychology, University of Montreal, Montreal, QC, Canada; Yale Child Study Center, Yale University, New Haven, CT 06520, USA; Department of Psychology, Goldsmiths, University of London; Department of Biology, McGill University, Quebec, QC, Canada H3A 1B1; School of Psychology, University of Auckland, Auckland 1010 New Zealand

## Abstract

Since Darwin^1^, researchers have proposed that human musicality evolved in a reproductive context in which males produce music to signal their mate quality to females. Sexually selected traits result from sex differences in the costs of high-quality signal production and high-fidelity signal detection^2^, leading to observable sexual dimorphisms across many species^3,4^. If musicality is a sexually selected trait in humans, males and females should then differ in their music perception ability, music production ability, or both. The evidence for this possibility is unclear, because previous reports of sex differences in human auditory perception are restricted in scope and inconsistent in direction^5–14^. Here, we report a test of music processing ability in >890,000 participants from 208 countries, over a main study and three replication studies. In contrast to other non-musical human traits^15–18^, and in contrast to music-related traits in non-human animals^19–22^, we found no consistent advantage in music perception or self-reported music production abilities for either sex. The sex differences we did observe in the main study (n = 554,300) were negligible (Cohen’s *d* range: 0.009-0.111), and Bayesian analyses of general music perception ability indicated evidence in favor of the null hypothesis of no sex difference (Bayes Factor = 0.6). This pattern of results held over the three replication studies. These results imply that the minds of men and women are largely similar when it comes to musicality, and so it is unlikely that sexual selection played any major role in the evolution of human musicality.

## Main Text

Music is ubiquitous across human societies, and there are numerous similarities in the forms and functions of musical styles across societies and cultures^23–31^. The biological and evolutionary roots of musicality, accordingly, have been debated at length^5,7,32–37^. One common hypothesis, originally proposed by Darwin^1^, and endorsed by many others^38–42^ is that music evolved via sexual selection as a credible signal of mate quality: males produce music to signal mate quality to females, while females assess potential mates based on their musical ability. This idea echoes credible signals of mate quality found in many non-human species, such as in the swamp sparrow (*Melospiza georgiana*), where females prefer male songs with rapid trills that are physically challenging to produce^43^; in birds-of-paradise (*Paradisaeidae*), where males evolved highly complex behavioral, morphological, and acoustic “courtship phenotypes”^44^; and in red-winged black birds (*Agelaius phoeniceus*), where females but not males perceive the difference between conspecific male calls and other species’ imitations of them^3^. Of course, humans are part of the animal kingdom, and in principle some of our vocalizations could operate similarly to these other species; indeed, human aesthetic judgments of animal calls align with those of numerous non-human species^45^.

Sexually selected traits demonstrate a reliable pattern of sexual dimorphism because the costs involved in reproduction differ across the sexes. In species in which females incur the larger cost for reproduction (e.g., larger gametes, internal fertilization, pregnancy, and lactation), females tend to be more discriminating of social signals such as courtship vocalizations because the cost of reproducing with a lower-quality mate is higher for females^2,3,46–49^. Indeed, female advantages in the sensory detection and discrimination of courtship signals have been documented in bird^19,20^, amphibian^50^, and mammalian species^51^. If music evolved in the context of sexual selection, human males and females should therefore differ in their music perception ability, music production ability, or both.

The evidence for this prediction is mixed^5,7–9,52,53^, and indeed the theoretical work does not clearly converge on a prediction for whether we might expect to see a female advantage in perception, a male advantage in production, or both. Some studies report a male advantage in musical analytic hearing abilities^10^; others report a female advantage in melodic recognition^12^ and auditory sensitivity^13^. Still others report no sex difference in pitch or beat perception abilities overall but possible differences in prevalence at the low extremes of these abilities^54^. Two studies that tested predictions related to the sexual selection theory of musicality have had mixed results: testosterone levels, as measured via saliva assays, were not predictive of men’s musical aptitude^11^, and in a large twin study, higher musical ability correlated with *lower* reproductive success^14^.

Further, several papers reporting evidence of sexual selection for musicality have been retracted or have documented failures to replicate: the claim that women are more attracted to men with apparent musical ability^55^ was retracted in 2020^56^; the claim that women are more attracted to higher quality dancing in men^57^, retracted in 2013^58^; and the claim that women prefer more complex music around ovulation^59^ failed to replicate^60^. Moreover, even in cases where a *bona fide* sex difference is found, such a difference could reflect sociocultural forces rather than biological forces^7^. This pattern has led to some skepticism regarding the sexual selection hypothesis for musicality evolution^5,7,53^.

Despite the skepticism, this hypothesis receives continued attention. For example, in a 2021 issue of 60 commentary articles on the evolution of music^53,61^, 12 commentaries refer to the hypothesis that music evolved under sexual selection. This hypothesis continues to be a focus in recent empirical work and books^36,62–68^.

Here, we employ a large-scale, citizen-science approach to examine sex differences in music perception. In a first study, we recruited 554,300 participants from 208 countries (see *Figure 1A* and *SI Table 1*) via https://themusiclab.org, a popular platform for gamified studies of auditory perception abilities^69^. Of these, *n* = 194,291 (34.5%) self-identified as “female”, and *n* = 360,009 (64%) self-identified as “male”. They completed three musical perception-based tests presented in a random order, measuring their i) mistuning perception^70^, ii) melodic discrimination^71^, and iii) beat alignment perception^72^. These tests cover core components of the human musicality phenotype^73^, and a previous use of these tests has shown that they are sensitive enough to detect sub-population-level effects, such as the effect of native-language experience on music processing abilities^74^. Participants reported demographic information, details of their musical training (if any), and listening habits (see *SI Table 2*). The participant sample was skewed somewhat male, possibly indicative of self-selection that could confound an estimation of sex differences; as such, we also report three replication studies to test the robustness of these findings. We begin with the results from the initial sample of 554,300 participants.

**Fig. 1.**
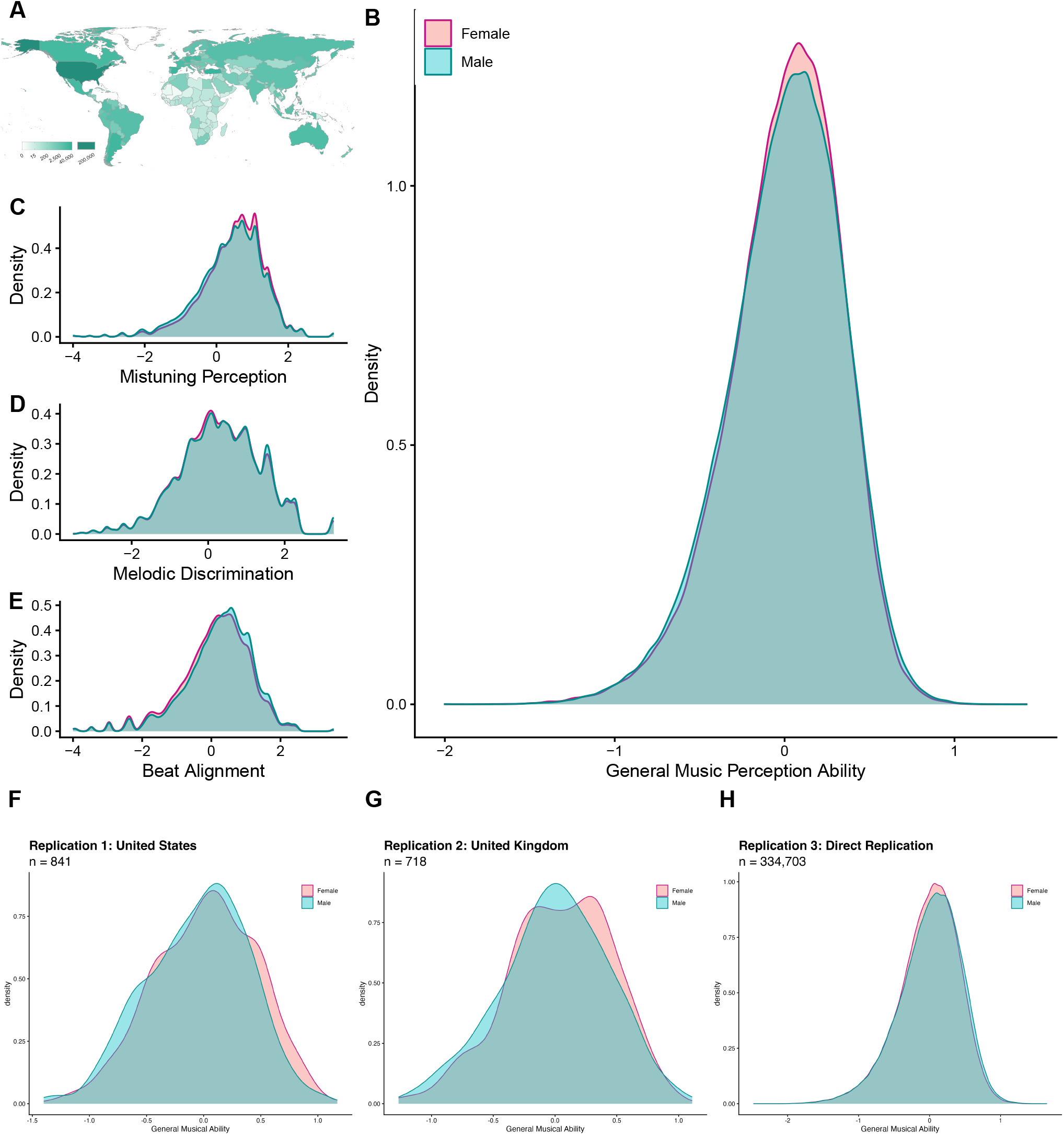
Negligibly small sex differences in music perception ability. A Geographic spread of participants who completed music perception tests. The shading of each country indicates the number of participants who self-reported that country as their location. B In an aggregate measure of general music perception ability (composed of measures C, D, E), there is a negligibly small difference between men and women’s scores. This pattern persisted in three replication studies: two in balanced samples of participants from F the United States (n = 447 women and n = 394 men), G the United Kingdom (n = 361 women and n = 357 men), and H a direct replication of the main study (n = 134936 women and n = 199767 men). The shaded areas represent kernel density estimations.

## Results

### Men and women differ minimally

We found statistically significant but negligibly sized sex differences in a measure of general music perception ability derived from the three tests, along with similarly negligible differences on each of the tests (p < 0.05 for each; *Figure 1B-E, Table 1*). Women, on average, scored higher than men on general music perception ability, but barely so (mean *±* SD; women: 0.002 *±* 0.331; men: -0.001*±* 0.34, Cohen’s *d* = 0.009). This difference is approximately 300 to 600 times smaller than average vocal acoustic differences between men and woman, a well-documented sexual dimorphism (Cohen’s *d* = 2.7 *−* 5.7^17^).

**Table 1.** Differences in male vs female music perception ability, quantified with a variety of effect size metrics. Note: Negative Cohen’s D values indicate that men’s scores are higher, and positive scores that women’s scores are higher.

| Task | Cohen's<br>D | %<br>Overlap | Probability<br>Superior | $\eta^2$ | $R^2$ | Bayes<br>Factor | Variance<br>Ratio<br>(m/f) |
| --- | --- | --- | --- | --- | --- | --- | --- |
| Mistuning Perception | 0.096 | 0.954 | 0.527 | 0.00205 | 0.00205 | 7.41e+244 | 1.101 |
| Melodic Discrimination | -0.020 | 0.934 | 0.494 | 0.00009 | 0.00009 | 4.49e+08 | 1.034 |
| Beat Alignment | -0.111 | 0.950 | 0.469 | 0.00280 | 0.00280 | Inf | 0.965 |
| General Music<br>Perception Ability | 0.009 | 0.980 | 0.503 | 0.00002 | 0.00002 | 6.07e-01 | 1.057 |

Further, a Bayesian approach to this analysis provided evidence leaning in favor of the null hypothesis of no true sex difference in general music perception ability (Bayes Factor = 0.6; *Table 1*). We also observe variance ratios (variance in male scores / variance in female scores) very close to 1 (range 0.965 - 1.101) (see SI).

The pattern of results on the individual perception tests was internally inconsistent. Women outperformed men in the mistuning perception test (women; 0.486 *±* 0.857; men: 0.402 *±* 0.9, Cohen’s d = 0.096), but men outperformed women on the beat alignment (women: m = 0.126 *±* 0.979; men: 0.234 *±* 0.962, Cohen’s d = -0.111) and melodic discrimination tests (women: 0.309 *±* 1.046; men: 0.33 *±* 1.064, Cohen’s d = -0.02). Bayesian approaches to these differences indicate that they are highly unlikely to be strictly zero (Bayes Factors >1000), despite their very small sizes and inconsistent directions. However, effects of such small magnitude would typically be considered to be null or very small effects^15,75,76^ (see *Table 1* for other effect size specifications).

This lack of clear sex differences cannot be explained by test insensitivity, as multiple other participant measures explained substantial variability in general music perception ability scores. For example, those participants who reported they had taken music lessons scored substantially higher than those who reported having never taken lessons (*d* = 0.68, i.e. 75 times larger than the main sex difference; *SI Figure 3*). Other work on a subset of these data report moderately-sized effects of language, with native speakers of tonal languages showing an advantage in melodic discrimination, of roughly half the size of the effect of having had music lessons^74^.

Furthermore, men and women’s self-assessed music perception and production skills were highly similar (see *Methods* for item details), and the relationship between self-reported factors and test scores expressed similarly between men and women. Similarly to the perception abilities we measured directly, men and women’s *self-assessed* musical listening abilities also differed minimally (answer scale 1-6: median answer 5 for both men and women; Wilcoxon test, W = 2.55e+10, *p* < .001, 95% CI [-0.00004, -0.00005], estimated higher score in men), comparable to the negligible difference in listening ability that we test directly here. Men and women also reported highly similar answers to questions pertaining to their ability to tap in time with a music beat (82.4% women answered “Yes” compared to 87.4% of men; logistic regression, *β* = 0.381, *z* = 23.073, *p* < .001, estimated higher score in men), their skill at making music (5 point scale: median answer 3 for both men and women; Wilcoxon test, W = 3.44e+10, *p* = 0.017, 95% CI [-0.00007, -0.00004], estimated higher score in men), and their ability to tell if they are singing out-of-tune (answer binary yes/no: 73.4% of women answered “Yes”, compared to 72.3% of men; logistic regression, *β* = -0.187, *z* = -16.798, *p* < .001, estimated higher score in women).

Participants’ self-assessed music production skill level was strongly related to performance on the music perception tests: those reporting they “have no skill” performed far worse on the listening tasks than those reporting being “experts” (Cohen’s *d* = 1.66; *Figure 2A*), and the magnitude of this difference between “no skill” and “expert” was similar when considering women and men separately (Cohen’s *d* = 1.53 and *d* = 1.73, respectively). Similarly, the degree of formal music training, as measured by self-reported participation in music theory classes, was highly related to performance on the tests for both men and women (*Figure 2B*); overall, those who self-reported that they had *“a lot of music theory training”* scored higher than those with *“no theory training”* (Cohen’s *d* = 0.9 globally; *d* = 0.914 for men, *d* = 0.881 for women).

**Fig. 2.**
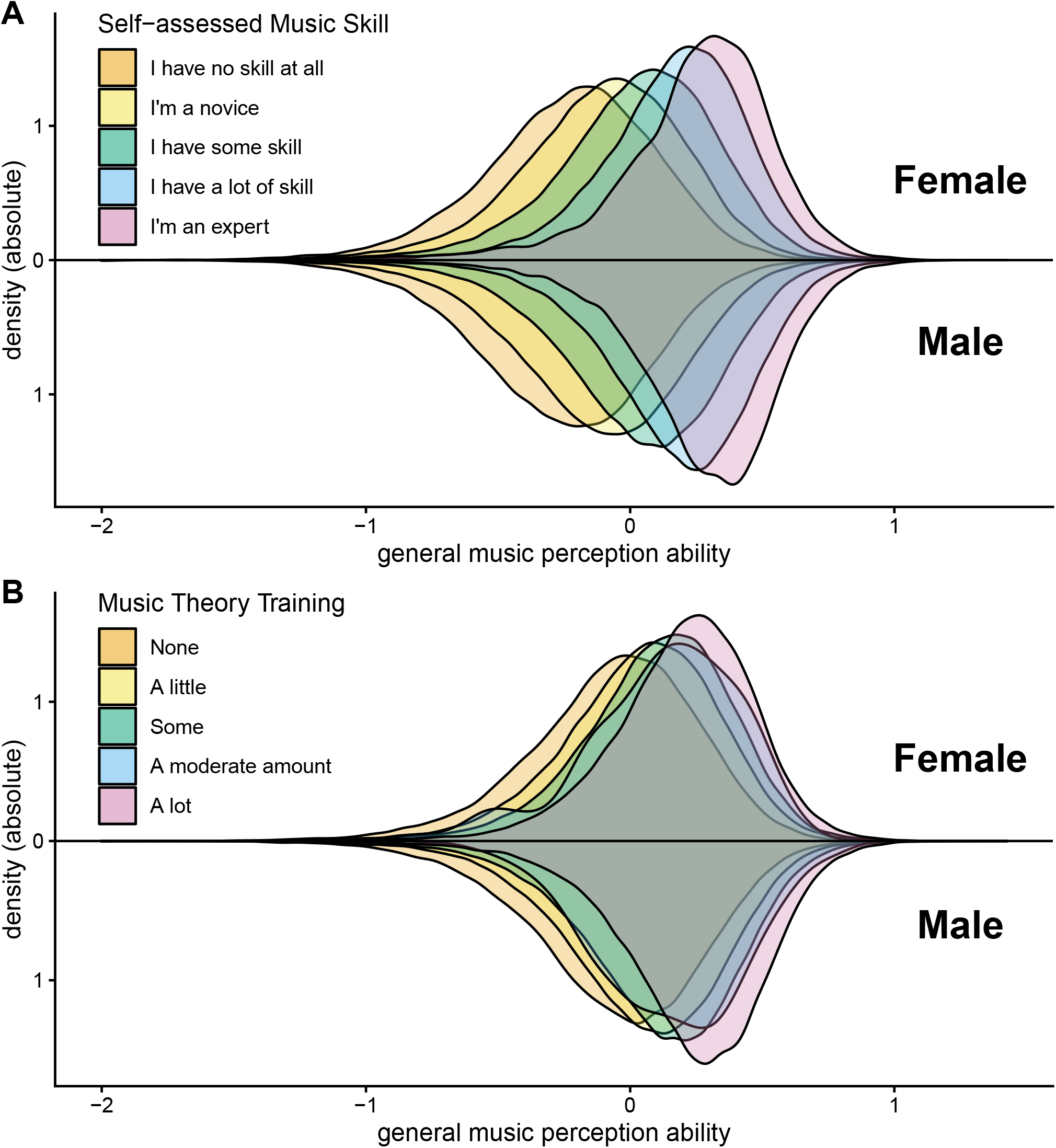
General music perception ability scores scale with self-reported musical skill and formal music education, but not with gender. As expected, general music perception ability scores differ much more along other participant covariates, such as self-estimated musical skill, than they differ by sex. The shaded areas represent kernel density estimations. The verbal prompt was *‘Think of your skill at making music (using a musical instrument or singing). How would you rate your own skill?’* The participant covariate of self-assessed musical skill captured much more variance in participants’ general music perception ability scores, compared to their sex (partial *η*^2^ =0.172 for self-assessed musical skill, relative to partial *η*^2^ =0.007 for sex). Cohen’s *d* values were all in the range of what would conventionally be described as medium effect sizes (Cohen’s *d* between ‘no skill’ and ‘novice’ = 0.396; between ‘novice’ and ‘some skill’ = 0.472; between ‘some skill’ and ‘a lot of skill’ = 0.486; between ‘a lot of skill’ and ‘expert’ = 0.339).

We also asked whether performance on the music perception tests might have been driven by participant covariates beyond their sex, and whether adjusting for these covariates changes the estimated magnitude of a sex difference. We used factor analysis to aggregate participant responses to questions concerning their musical skill, training, and listening habits (for details about factors, see *Methods, & Table 4*). We then repeated the main analyses while adjusting for participants’ age, education, age of start of music training (if any), and the above factors. The inclusion of these covariates did not substantially alter any effect sizes of sex differences (*SI Table 2*). Further analyses using structural equations modelling approaches yielded comparable results (see *Analysis & SI Figure 1*).

### Negligible sex differences hold across three replication studies

The male skew of the initial participant sample (64.9% male) could indicate a difference in how men and women self-selected into the study, which may have created a confound to our estimation of sex differences. Indeed, anecdotally, we noted that large spikes in website traffic temporally coincided with social media attention from YouTube, Twitch, and Reddit, the latter platform having a notably male-skewed audience^77^. We tested whether a different recruitment strategy would lead to different conclusions about sex differences on the music perception tests. We recruited additional participants in the United States (Replication 1) and the United Kingdom (Replication 2), each balanced on gender and stratified by age and education (see Methods, Table 3), who then took the same tests as the participants in the main study. Additionally, in the time since the original data analysis, the gamified test continued to accumulate participants, after the initial wave of social media attention to the game. We pre-registered the analysis of the replication data (Replication 3; repregistration available at https://osf.io/aczdv/overview?view_only=06b3e339de4a4621b8b9f6bb3de4f7d6).

If there had been a large self-selection effect in the first, main sample that confounded the original estimate of sex differences in music perception ability, the estimated sex differences in these replication sample would be substantially different to the original estimate. This was not the case. Just as in the original citizen-science sample, the sample of 841 US participants that were externally recruited and demographically balanced (Replication 1) displayed a pattern of small sex differences in the three tests and aggregate general musical perception ability score (0.059 < |Cohen’s *d*| < 0.18), with inconsistent directions and low Bayes Factors (*Figure 1F, Table 2*).

**Table 2.**
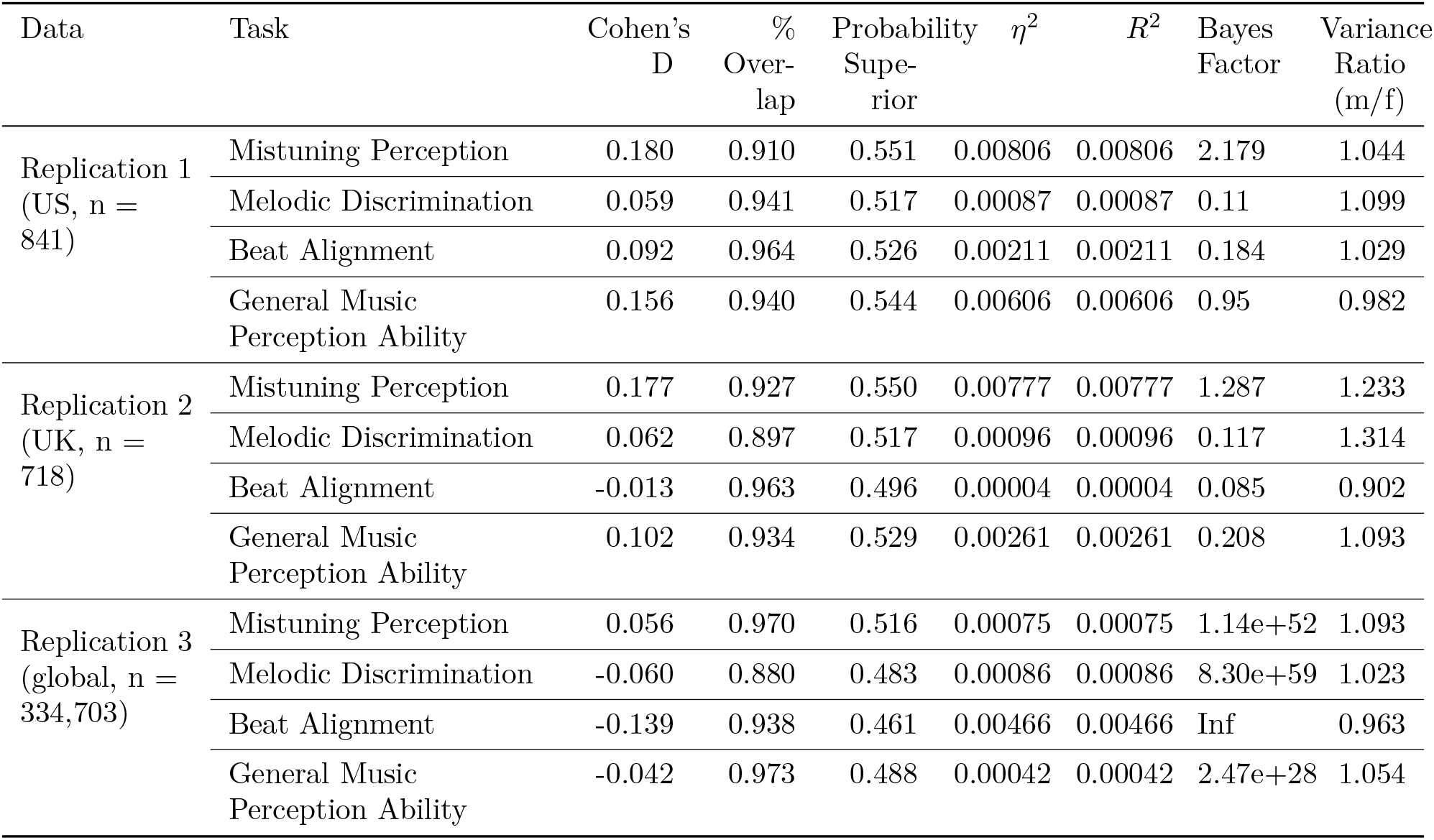
Effect sizes across three replication studies. Across three replication studies, we continue to see a pattern of functionally negligible sex differences in music perception abilities. Negative Cohen’s D values indicate that men’s scores are higher, and positive scores that women’s scores are higher.

| Data | Task | Cohen's<br>D | %<br>Over-<br>lap | Probability<br>Supe-<br>rior | $\eta^2$ | $R^2$ | Bayes<br>Factor | Variance<br>Ratio<br>(m/f) |
| --- | --- | --- | --- | --- | --- | --- | --- | --- |
| Replication 1<br>(US, n =<br>841) | Mistuning Perception | 0.180 | 0.910 | 0.551 | 0.00806 | 0.00806 | 2.179 | 1.044 |
|  | Melodic Discrimination | 0.059 | 0.941 | 0.517 | 0.00087 | 0.00087 | 0.11 | 1.099 |
|  | Beat Alignment | 0.092 | 0.964 | 0.526 | 0.00211 | 0.00211 | 0.184 | 1.029 |
|  | General Music<br>Perception Ability | 0.156 | 0.940 | 0.544 | 0.00606 | 0.00606 | 0.95 | 0.982 |
| Replication 2<br>(UK, n =<br>718) | Mistuning Perception | 0.177 | 0.927 | 0.550 | 0.00777 | 0.00777 | 1.287 | 1.233 |
|  | Melodic Discrimination | 0.062 | 0.897 | 0.517 | 0.00096 | 0.00096 | 0.117 | 1.314 |
|  | Beat Alignment | -0.013 | 0.963 | 0.496 | 0.00004 | 0.00004 | 0.085 | 0.902 |
|  | General Music<br>Perception Ability | 0.102 | 0.934 | 0.529 | 0.00261 | 0.00261 | 0.208 | 1.093 |
| Replication 3<br>(global, n =<br>334,703) | Mistuning Perception | 0.056 | 0.970 | 0.516 | 0.00075 | 0.00075 | 1.14e+52 | 1.093 |
|  | Melodic Discrimination | -0.060 | 0.880 | 0.483 | 0.00086 | 0.00086 | 8.30e+59 | 1.023 |
|  | Beat Alignment | -0.139 | 0.938 | 0.461 | 0.00466 | 0.00466 | Inf | 0.963 |
|  | General Music<br>Perception Ability | -0.042 | 0.973 | 0.488 | 0.00042 | 0.00042 | 2.47e+28 | 1.054 |

Replication 2 afforded us a second country replication with this age-, gender-, and education-stratified participant sampling method. Here, too, we found that these 718 United Kingdom male and female participants showed small differences in the three tests and aggregate general music perception ability (0.013 < |Cohen’s *d*| < 0.177; *Figure 1G, Table 2*), consistent with the original study, and Replication 1.

In Replication 3’s additional 334,703 participants (n = 134,936 women, n = 199,767 men), we observed a pattern of small sex differences in inconsistent directions (see *Figure 1, Table 1*), in line with the main finding and Replications 1 and 2. The sex difference in aggregate music perception ability was estimated to be d = -0.042, (men scoring higher), aggregating minimal differences for mistuning perception (Cohen’s d = 0.056, women scoring higher), melodic discrimination (Cohen’s d = -0.06, men scoring higher), and beat perception (Cohen’s d = -0.139, men scoring higher).

### Age, education, musical training, environmental noise, and country explain more score variance than sex

To contextualize how small the influence participant gender was on music perception abilities, we took the largest effect size in any of the samples we tested - the Cohen’s *d* of 0.18 in Replication 1 US participants’ scores on mistuning perception - and fit an additional linear model that computed this effect size after accounting for participant age, education, musical training, and noisiness of the environment in which they completed the task (i.e. a 5-point scale ranging from *“I am in a very quiet place”* to *“I am in a very noisy place”*). In the simple model that fits test scores simply as a factor of gender, this Cohen’s d of 0.18 corresponds to an *η*^2^ = 0.008. In a model with the above covariates, this drops slightly to 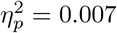, which is smaller than the variance explained by any of the other individual factors: we observed larger partial *η*s for self-reported noisiness of the space in which the participant completed the test 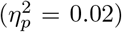, age 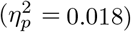, education 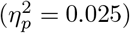, and years of music training 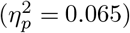 . Even in the largest sex difference we observed across four participant samples (main study and three replications), sex accounted for a vanishingly small amount of variance.

Last, we asked how uniform this sex difference was, globally. We re-calculated the magnitude of the sex difference on general music perception ability *within each country*. We pooled the main study and direct replication data (Replication 3; for a total n = 889,003 from 210 countries), and filtered the data to retain countries with large sample sizes (*n* ≥ 1000) to build well-sampled distributions of test scores. Of these 70 countries, we computed per-country average sex differences in general music perception ability, and Cohen’s d effect sizes for these differences. Despite that listeners from different countries differ in test performance (possibly due to differences in familiarity to Western music styles in the test stimuli; see below), the magnitude of estimated sex differences in general musical perception ability appeared highly similar across countries, with the distribution of average test score differences overlapping with 0 (mean score difference = -0.009, sd = 0.044, *t* = -1.631, *p* = 0.107; mean Cohen’s d = -0.025; *Figure 3*), and orders of magnitude smaller than cross-cultural estimations of average sex differences in known sexual dimorphisms, such as height^16^ or vocal acoustics^17^ (*Figure 3B*).

**Fig. 3.**
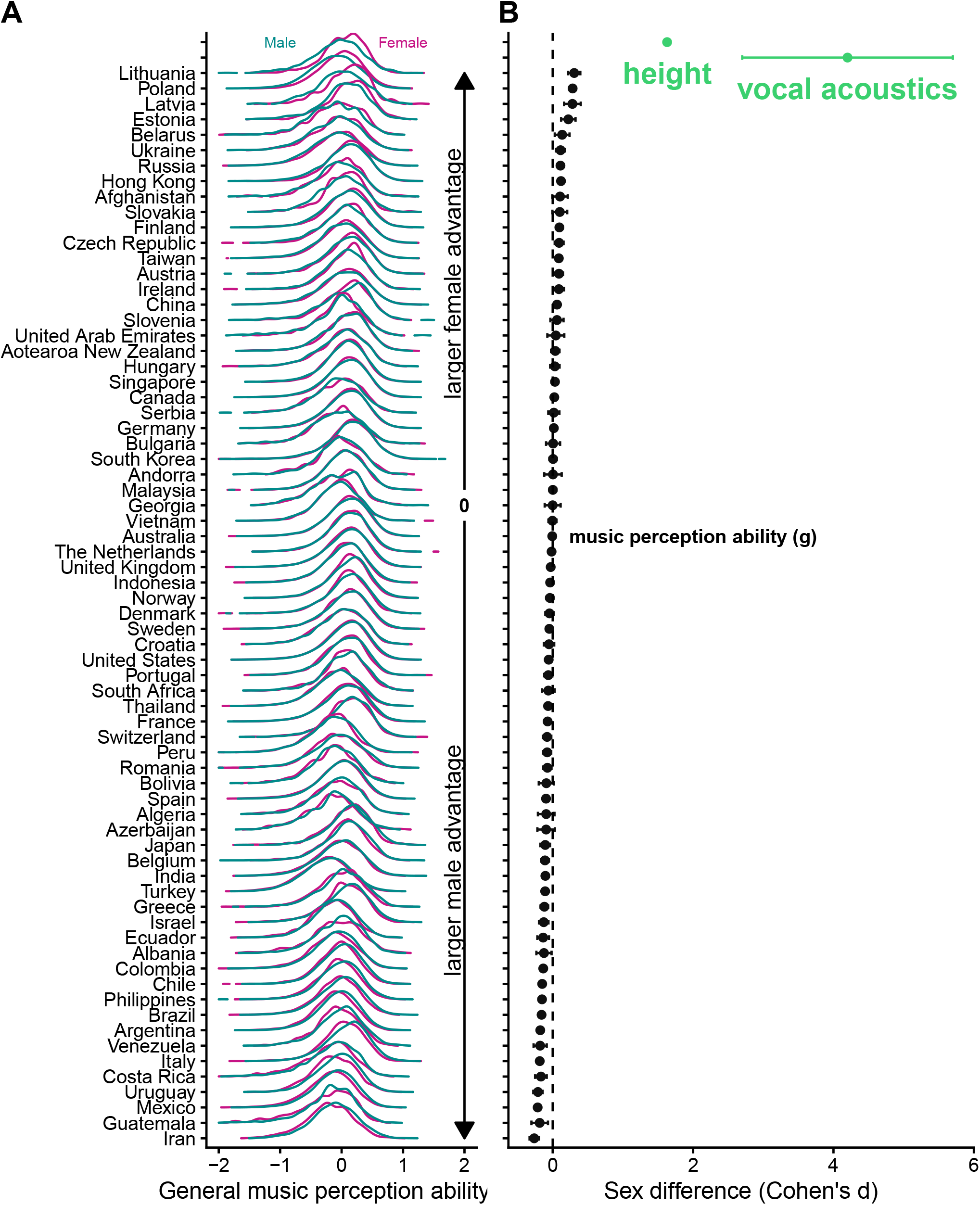
Per-country variation in sex differences clusters around 0. Only displaying countries with n > 1000 participants. A Per-country distributions of male and female participants’ scores on general music perception ability, visualized as density plots, ordered by direction and magnitude of sex difference in mean score. B For these same countries, equivalent Cohen’s d that describe the sex difference in these distributions, in the same order as in A, with whiskers marking 95% confidence intervals. For voice differences, the range marked are minimum and maximum Cohen’s ds on various acoustic features (from Puts et al., 2011). For height difference, only a point estimate was available (from Lippa et al., 2009).

Moreover, the effect of country was orders of magnitude larger than the observed sex differences. We ran linear models that fit test scores using only sex information, or sex and country information. The latter model had better fit (*R*^2^ of 0.000044 and 0.067, respectively), and in the latter model, the partial *η* of sex 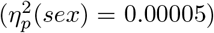 was several orders of magnitude smaller than that of country 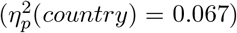 .

## Discussion

Together, these findings provide no evidence for substantive sex differences in human music perception. Not only did the most general test of sex differences only find weak evidence for a sex difference (i.e. a negligible effect size, and a low Bayes Factor of 0.6), but the reliable sex differences we *did* find were of a very small size and of inconsistent direction. This pattern held across three replication studies; in US and UK participant samples, where recruitment did not rely on organic social media “word of mouth”; and again in a large-scale direct replication.

Those few reliable differences do not seem to support any claim of sexual selection, for two main reasons. First, human sex differences in traits argued to result from sexual selection tend to be large in size; some examples include height (Cohen’s *d* = 1.63)^16^, vocal acoustic features (Cohen’s *d* = 2.7 - 5.7)^17^, and rates of engagement in physical aggression (Cohen’s *d* = 1.11)^15^. These effects dwarf the sex differences we found on the three music tests. The small sex differences that we do observe appear easily overridden by factors like age, country, education, and simple self-reported noisiness of the environment in which one does a music listening test.

Second, large sex differences in perception have been observed in a variety of non-human animals that *do* use acoustic signals for courtship and sexual behavior. Female black-capped chickadees (*Poecile atricapillus*) outperform males on the discrimination of pitch ratios 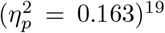, and are significantly faster than males at discriminating between socially dominant and subordinate male songs (Cohen’s *d* = 2.854)^20^ ; and only female Bengalese finches showed recognition of new males’ songs (as seen in heart rate habituation)^78^. Because in these species males and females often have categorically different behavioral responses to song, studies directly comparable to our own—where both males and females are presented with the same stimuli and assessed on the same behavioral assay—are sparse in the literature, and so there is little consensus as to whether these sex differences are strictly perceptual and/or motivational. While the exact mechanisms underlying these sex differences in non-humans animals’ responses to these courtship signals is unclear, the magnitude of non-human animals’ sex differences in these behavioral responses equally dwarf the human sex differences we found on the three music tests.

The lack of sex differences in music perception casts doubt on the mate quality hypothesis, but does not rule it out entirely. The ability to produce music requires complex sensory processing and sensorimotor integration; one can imagine unusual co-evolutionary dynamics at work, where music production is substantively dimorphic but music perception is not. We find this unlikely: in our data, self-reported music production skill was highly correlated with music perception ability in both males and females, suggesting there should be similarly small sex differences in music production skills. This is consistent with a large twin study finding no sex difference in singing abilities^79^, and other work finding associations between music perception and production abilities^70–72,80^. While self-reports of music production ability are coarse measures relative to structured assessments of music production ability, we predict that future research using such tests will find comparably poor evidence for sexual dimorphism in musicality, and, therefore, little support for the sexual selection theory of the evolution of musicality.

## Methods

This research was approved by the Harvard University Committee on the Use of Human Subjects (protocol 2000033433) and Yale University’s Human Research Protection Program (protocol 2000033433). Data were collected via an online experiment hosted at https://www.themusiclab.org/quizzes/miq which was advertised as a “Test Your Musical IQ” game.

### Participants

#### Main Study

Recruitment was driven by organic social media sharing, which lead to the gamified experiment reaching 562,853 participants from 208 countries. The first round of data collection was gathered between 22 Nov 2019 and 14 Dec 2020. Included in this dataset are only individuals who self-reported that they had not taken the test before and did not have a hearing impairment. Of these, n = 194,291 (34.5%) self-identified as “female”, n = 360,009 (64%) self-identified as “male” and n = 7,521 (1.3%) indicated “other” for their gender. Because sexual selection theory does not make explicit predictions with regards to how non-binary participants may differ from men and women, we only used data from participants who self-identified as either male or female. While individuals from over 200 countries participated in the test, the majority of participants were from North America (173,690, or 30.9%) and Europe (222,337, or 39.5%). Further information about participants’ age and educational achievement is detailed in Table 3 and SI Table 1.

**Table 3.**
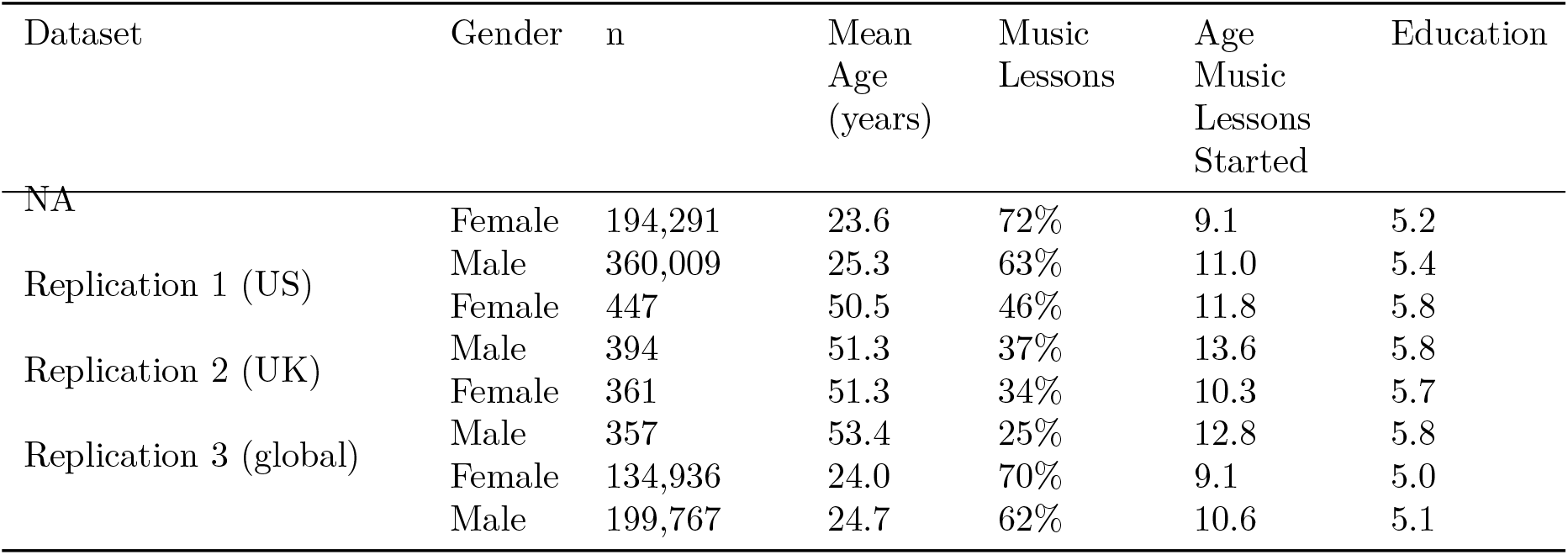
Participants. Participant information, broken down by self-reported gender, age, education, and music lessons. ‘Music Lessons’ was a binary question asking whether participants had ever had music lessons or not. ‘Age Music Lessons Started’ asked, of those participants that had had lessons, when they started. ‘Education’ answers ranged from ‘Some elementary school’ (scored 1) to ‘Completed graduate school’ (scored 8); the average education levels in these populations are between ‘Some undergrad’ (scored 5) and ‘completed undergrad degree’ (scored 6).

**Table 4.**
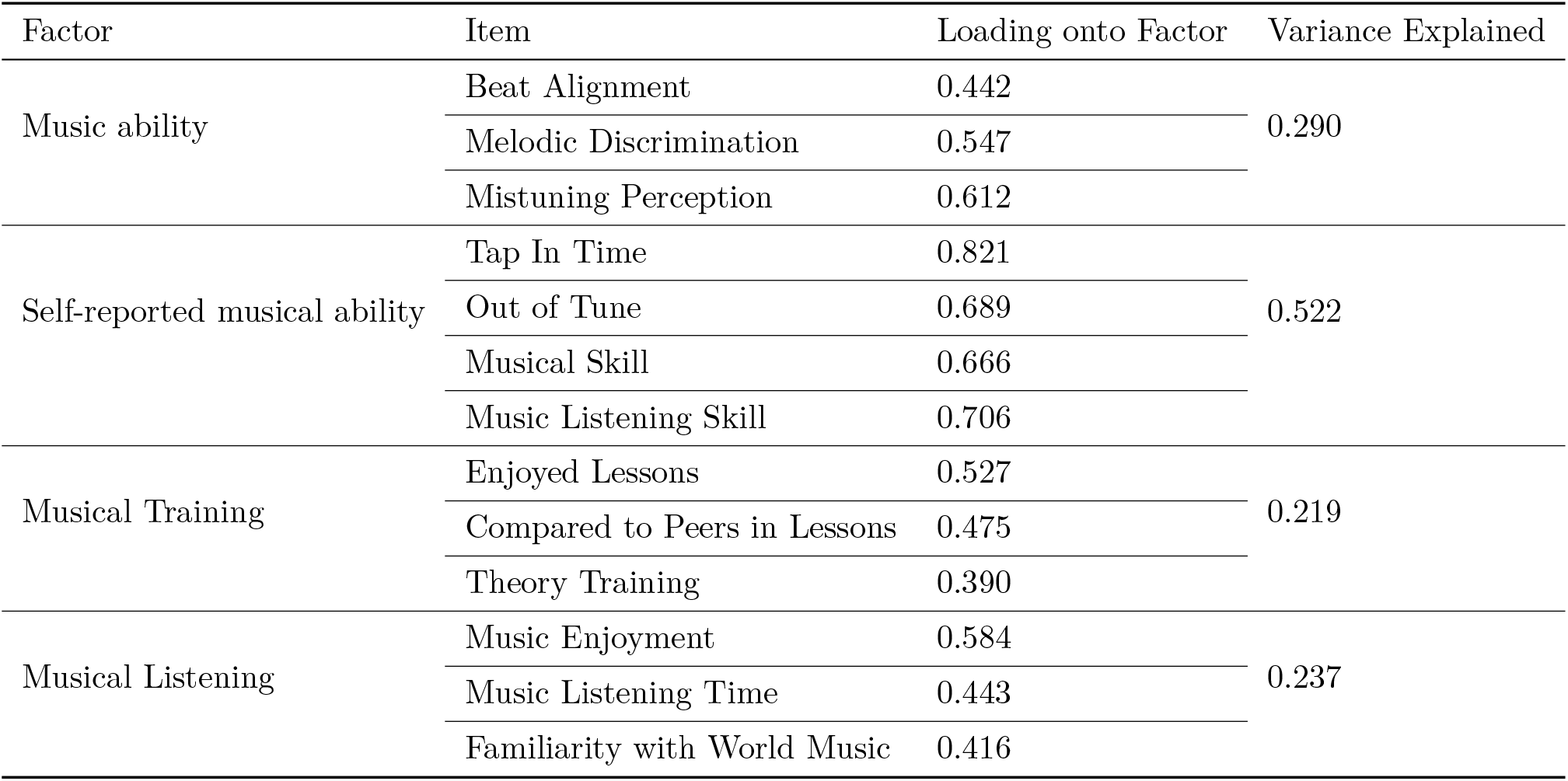
Factor Analysis. Aggregation of related items into factors, and individual items’ loadings onto their factor. Factor analysis was applied to data from the main study.

| Factor | Item | Loading onto Factor | Variance Explained |
| --- | --- | --- | --- |
| Music ability | Beat Alignment | 0.442 | 0.290 |
|  | Melodic Discrimination | 0.547 |  |
|  | Mistuning Perception | 0.612 |  |
| Self-reported musical ability | Tap In Time | 0.821 | 0.522 |
|  | Out of Tune | 0.689 |  |
|  | Musical Skill | 0.666 |  |
|  | Music Listening Skill | 0.706 |  |
| Musical Training | Enjoyed Lessons | 0.527 | 0.219 |
|  | Compared to Peers in Lessons | 0.475 |  |
|  | Theory Training | 0.390 |  |
| Musical Listening | Music Enjoyment | 0.584 | 0.237 |
|  | Music Listening Time | 0.443 |  |
|  | Familiarity with World Music | 0.416 |  |

#### Replication Studies 1 & 2

We worked with SoundOut, a third-party collaborator that specialises in commercial music testing, to build participant samples for the US and the UK, with each sample balanced on numbers of men and women, and men and women in each sample balanced on age and education (see *Table 3*). Their responses to the demographic portion of our study indicated that they were also balanced on musical education. Of the target n = 1000 per country, n = 841 (447 female, 394 male) from the US and n = 718 (361 female, 357 male) from the UK provided complete data, reported that they had not taken the test before, and had no hearing impairments.

#### Replication Study 3

In the time since the original analysis of the data and conducting the above replication studies, the original test hosted on https://www.themusiclab.org/ continued to organically accumulate participants. This allowed us to conduct a third replication study, this one a direct replication of the main finding. These data (collected 14 Dec 2020 to 09 March 2026) form a sample of an additional n = 334,703 from 206 countries. These participants completed the same tests as the original sample and Replication Studies 1 and 2, and were identically filtered on age (retaining those who self-reported an age between 7-100), having self-reported never taken the test before, and self-reported having no hearing impairments.

#### Measures of music-perceptual abilities

Participants completed three musical perception-based tests, each of which were based on prior work. These tests used an explanatory item response theory model and were adaptive. Each of the three tests was 15 items long, and the adaptive nature of the tests allows the procedure to select items dynamically such that item difficulty is matched to estimated participant ability for each trial. The exact 15 items (per test) that participants heard therefore varied according to their test performance. Validation for item difficulty, algorithms for item difficulty selection, and scoring calculations are reported in the original papers that validated these tasks. Final participant scores, per test, are computed according to the underlying item response theory model and have a theoretical range from -4 to 4, with higher scores indicating greater ability. These tests measures participants’ abilities in the following domains:

i. Melodic discrimination^71^: Participants listen to three different versions of the same melody. All three versions are played at different absolute pitch levels, with subsequent versions always transposed a semitone higher compared to the previous version. Two versions are identical in their interval structure but one of the three versions differs in one note. Differences can be in melodic interval, contour or tonality or any combination of the three factors. The participant’s task is to identify the odd-one-out. Melodies were synthesized using MIDI with a piano timbre; they varied in length, ranging between 6-16 notes depending on difficulty level; and were all presented at 120 beats per minute.
ii. Mistuning perception^70^: Participants are given two nearly identical short excerpts of vocal music. In one excerpt the vocal track was pitch-shifted against the instrumental backtrack; the participant’s task is to indicate in which of the two excerpts the singer sounds more out of tune. Of the 45 items in this test, 26 had male vocalists. For each item, de-tuned versions were generated with pitch-shifts ranging from 10-100 cents (in 5 cent increments; 100 cents corresponds to one semitone), thus generating item pairs of different difficulty levels.

Beat alignment^72^: Participants listen to the same clip of music twice, with an overlaid beep track. In one version the beep track is perfectly aligned with the musical beat of the music, and in the other version the beep track is slightly shifted in time. The beep track was offset from the musical beat by a constant proportion of the musical beat, that ranged 0 < P ≤ 0.5. A deviation if 0.5 indicates a maximum deviation of being half a beat misaligned with the musical beat, thus creating the easiest items to discriminate against the clip with the perfectly aligned beep track. The beep track could be misaligned from the musical beat by being ahead of or behind the musical beat. The participant’s task is to identify in which of the two clips the beep track is best aligned with the beat of the music.

The music consisted of excerpts of Western popular music, i.e., music in a style that would be familiar to listeners acculturated to Western music, but not songs that they were likely to have heard before; all stimulus materials were sourced from stock libraries containing only songs that have not been commercially released. While our tests consisted only of musical excerpts in the Western musical tradition, because men and women from all geographic regions participated in the study, this test remains useful for quantifying sex differences around the world; any culture-driven differences in performance would apply equally to both men and women, and therefore are unlikely to confound the presently reported findings.

### Procedure

Participants completed the experiment on computers, smartphones, or tablets. Participants were first asked to report their age, country, self-identified gender, native language, and whether they had a hearing impairment. They were also asked about the noisiness of the environment in which they were at the time of the experiment, whether they were wearing headphones, and if they had taken the test before. Participants then completed the three music perception tests, in a randomized order. After the tests, they were asked a number of questions concerning their musical skill and experience (see *SI Table 3*).

## Analysis

### Data pre-processing

Of the >2 million people who had begun the online experiment at https://www.themusiclab.org/miq between 22 Nov 2019 and 14 Dec 2020 (main study), we considered only those who had completed all three musical perception tests, only analyzed the data from participants who self-reportedly had not taken the test before, and excluded participants who reported an age younger than 7 or older than 100. Applying these filters resulted in the main study’s participant sample of n = 562,853. Questions that had written response labels were re-coded into binary or ordinal variables (see *SI Table 3*); these items formed conceptually meaningful groups: items pertaining to *i)* musical perception ability (ie. the three musical perception tests), *ii)* self-reported musical ability, *iii)* formal musical training, and *iv)* extent, intensity, and breadth of musical listening. These same data pre-processing steps were to the three replication datasets.

### Factor analysis to extract composite scores

For each of the above groups of measures, information from their constituent sources was aggregated by factor analysis. For each of these factor analyses, the data from the main study was randomly split into two equal sized subsamples. An exploratory minimum residual factor analysis was run on one subsample, with the correlation matrix of the variables belonging to the same conceptual group. Questions differed in their response options and, by extension, the measurement level of corresponding variables. In these cases a mixed-type correlation matrix was computed employing polychoric, tetrachoric, biserial, point-biserial or Pearson correlations as appropriate for each pair of questions. The resulting correlation matrix was then subjected to a factor analysis. In all cases only a single factor was specified and only variables that loaded with a minimum of 0.3 were retained in the factor model. For example, the variable for whether participants had had music lessons, which participants answer with a binary yes/no, was pruned due to being too coarse of a measure to capture variation in test scores; the “yes” answer to this item pools participants who only briefly had lessons with those who reported decades of training. Other factors pertaining to music training (i.e. whether participants enjoyed lessons, their self-assessment of skill relative to peers, and amount of music theory training, including “none”) were found to load sufficiently onto the Musical Training factor. The second subsample was used for a confirmatory factor analysis and the factor structure was confirmed by inspecting measures of absolute model fit. On the condition that fit measures were satisfactory, the confirmatory factor model was then computed a second time, using the entire sample of participants, and factor scores were computed using only data from those participants with complete data on all the variables included in the factor model. We then used the Empirical Bayes Modal approach for computing factor scores from the confirmatory model. Table 3 details each factor’s individual items, their loading on the factor, and how much variance among the indicators is explained by the single factor. For all factors, the confirmatory fit measures indicated a good model fit (General music perception ability RMSEA < .001, SRMR, < .001; self-reported musical ability RMSEA = 0.029, SRMR = 0.037; self-reported musical training RMSEA < .001, SRMR, < .001; self-reported musical listening RMSEA < .001, SRMR, < .001).

### Estimating sex differences in music perception

To test whether there were any sex differences in music perception in the main study, after controlling for the above participant covariates, we tested for sex differences using a sequence of different analysis approaches.

First, a general additive model (GAM) was fit to each dependent variable, i.e., the three individual ability tests (melodic discrimination, mistuning perception, and beat alignment), and the aggregate factor score of general music perception ability. This allowed us to calculate means and variances of these variables and test whether they differed between men and women. Men and women’s distributions on each variable were plotted to visualize the extent of their overlap (*Figure 1*). On the basis of these data, we also calculated the probability superior (the probability that a randomly selected woman would have a higher score than a randomly selected man), the *η*^2^ of how much variance sex explains in the models, the *R*^2^ of the models, and the variance ratios (variance men / variance women) (*Table 1*). As a measure of effect size of a potential sex difference, we calculated Cohen’s *d* as a parametric effect size, and percentage overlap of the two distributions as a non-parametric effect size (*Table 1*).

Secondly, in order to further test whether men and women differed on this latent variable “general music perception ability” that summarizes the three musical perception tests, we computed structural equation models (SEMs) using the factor scores on the three perceptual tests as indicators of a latent variable general music perception ability (*g*) (see *SI Figure 1*). We constrained the intercepts and the loadings of the three tests on *g* to be equal across men and women, tested the difference in the latent means of *g* for significance, and computed Cohen’s *d* effect sizes. This model estimated that the difference in *g* between men and women was 0.009, with a pooled variance of 0.552. This corresponds to an effect size of *d* = 0.016, which is comparable to the above estimates derived from the GAM.

### Controlling for participant covariates

To test whether this magnitude of difference in *g* may have been confounded by other participant variables in the main study, the above SEM was extended to include other participant variables that may have plausibly been influencing their test performance - specifically participants’ age, education, age at which they started music lessons (if any), musical training, and musical listening. This SEM approach additionally allows us to model the correlational structure among the covariates (see *SI Figure 1*). When taking covariate effects into account, the effect size increased slightly: the difference in *g* after covariate adjustment was 0.051, with a pooled variance of 0.47. This corresponds to an effect size of *d* = 0.108. This may reflect the influence of slight group differences on the above covariates.

### Testing whether differences were robust across covariate subgroups

Lastly, we tested whether the very small sex differences observed above, in the main study, were robust across different combinations of participants’ covariates (namely: age, education, start age of music lessons, and factors musical training and musical listening). To do this, we used a tree model based on recursive partitioning. The tree model splits covariates into subgroups for which the coefficient for sex in a linear model changes significantly. Given our large sample size, we set a significance threshold of p < .001 and required a minimum subgroup to have at least 1% of the total sample size. Tree models were automatically pruned using the Bayesian Information Criterion. We computed the range of the effect sizes (Cohen’s *d*) across subgroups for the tree model of each dependent variable.

Most of these tree models formed subgroups based on participants’ amount of musical training and start age of music lessons. The range of effect sizes for sex across these tree model subgroups ranged from Cohen’s *d* = 0.005 to 0.31 (see *SI Table 4* for ranges per test), this higher end representing effect sizes that are conventionally considered small effects (i.e. 0.2 < *d* < 0.5)^81^. Across these tree models, most models estimated a sex difference of Cohen’s *d* < 0.2, and lower and upper bound for these estimates overlapped with 0 for mistuning perception, beat alignment, and general music perception ability. For melodic discrimination, the range of sex differences across these subgroups was [0.017; 0.179], indicating a very small female advantage. Even when controlling for other participant variables, sex differences on the three tests, and general musical perception ability, are very small.

These tree models reveal that subgroups constructed from these other variables reveal slightly larger effect sizes for sex than those estimated in the full distribution. These can be explained by the covariate musical training. Musical training is positively related to musical ability, and females participants reported having higher musical training on average. Hence, when data are conditioned on musical training (i.e. males and females are made statistically equal in terms of training), the estimated sex difference effect size slightly increased, with a small male advantage in most subgroups. This appears to be a version of Simpson’s paradox, where there are (almost) no differences in the full sample, but slightly stronger differences emerge for (almost) all subgroups analysed. To contextualize this apparent indication that conditioning on musical training increased the estimated effect size, we reran the main analysis of sex differences on the subset of participants who reported no musical training (i.e. participants who were, in fact, matched on musical training). Among participants that reported no musical training, we observed a sex difference on general music perception ability of magnitude Cohen’s d = -0.099 (negative score indicating a male advantage; mistuning perception Cohen’s d = 0.014; melodic discrimination Cohen’s d = 0.014; beat alignment Cohen’s d = -0.17; see *SI Figure 2* for male and female distributions on general music perception ability, for “yes” and “no” music training). Here, too, all estimated sex differences fall under the range of what are considered very small, and inconsistently estimating an advantage for men or women.

### Testing for differences in self-assessed music skills

On the four questions that asked about the main study’s participants’ self-reported musical ability (see *SI Table 3*), men and women’s responses to them differed minimally. We tested for differences in mean responses using logistic regression where the answers were binary (Yes/No questions), and Wilcoxon tests where answers were ordinal (i.e. answers on a scale of 1-5 or 1-6).

i. “Can you tap in time with a musical beat?” (answer binary yes/no).
ii. “How good do you think your musical listening skills are (things like remembering melodies, hearing out of tune notes, or hearing a beat that is out of sync with the music)?” (6-point scale).
iii. “Think of your skill at making music (using a musical instrument or singing). How would you rate your own skill?” (5-point scale).
iv. “When you are singing, can you tell if you are out-of-tune or off-key?” (answer binary yes/no).

## End notes

### Data, code, and materials availability

A fully reproducible manuscript, with data, analysis code, and visualizations is available at https://github.com/themusiclab/sex-differences. Readers may participate in the perception test by visiting https://themusiclab.org/quizzes/miq.

## Acknowledgments

This research was supported by the Harvard Data Science Initiative (S.A.M.), the National Institutes of Health Director’s Early Independence Award DP5OD024566 (S.A.M), the Royal Society of New Zealand Te Apārangi Rutherford Discovery Fellowship RDF-UOA2103 (S.A.M.), Natural Sciences and Engineering Research Council of Canada, Grant/Award Number: 05016 (J.T.S.), Fonds de Recherche du Québec Nature et Technologies, Grant/Award Number: PR-299652 (S.C.W.), an Anneliese Maier research prize awarded by the Humboldt Foundation (D.M.), a doctoral training grant from Fonds de Recherche du Québec Nature et Technologies https://doi.org/10.69777/348674 (M.B.), and the Center for Research on Brain, Language, and Music (M.B.). We thank the participants, Christopher Sturdy, and members of The Music Lab and The Peretz Lab for feedback, Courtney Hilton and Peter Harrison for technical assistance.

## Author contributions

Conception S.A.M., D.M.; experimental design and implementation, S.A.M.; participant recruitment, data management, and data processing, S.A.M., D.M., M.B.; analysis and visualization, M.B., D.M.; writing, M.B., D.M., I.P., S.C.W., J.T.S., S.A.M.

## Supplementary Information

### Supplementary Text: Is there greater male variability?

The “Greater Male Variability Hypothesis” argues that some dimorphisms manifest as greater variability - not greater averages - in males than in females (e.g., “more idiots, more geniuses”^35^). Significant (p < 0.05 for each) but small gender differences in variances were observed, but like the differences in mean performance found in individual tests, they were of internally inconsistent directions: variances were significantly larger for men in the mistuning perception (men:women variance ratio: 1.101) and melodic discrimination tests (men:women variance ratio: 1.034), and larger for women in the beat alignment test (men:women variance ratio: 0.965; see *Tables 1, 2* for variance ratios for all tests, in the main and replication studies). The variance ratios all being close to 1, however, indicates highly similar variances between men and women^75^. Thus, we find no clear evidence in support of higher male variability in music perception ability.

### Supplementary Tables

**SI Table 1.** Participant information for the main study, broken down by self-reported gender, age, and education. Percentages per row do not add up to 100 because not all participants opted to report education information.

| Gender | n | Age<br>(years,<br>mean) | Age (SD) | Highest education completed |  |  |  |
| --- | --- | --- | --- | --- | --- | --- | --- |
|  |  |  |  | High Schoool | Some Undergrad | Undergrad | Postgrad |
| Female | 194,291 | 23.6 | 9.6 | 30824 (16%) | 32317 (17%) | 45859 (24%) | 35765 (18%) |
| Male | 360,009 | 25.3 | 9.5 | 64320 (18%) | 66161 (18%) | 88386 (25%) | 72823 (20%) |
| Other | 7,521 | 23.0 | 9.7 | 1362 (18%) | 1550 (21%) | 1662 (22%) | 854 (11%) |
| NA | 1,032 | 25.7 | 9.0 | 166 (16%) | 152 (15%) | 206 (20%) | 189 (18%) |

**SI Table 2.** Full model, gender and covariates. The estimated gender effect remains of a comparable magnitude when other participant covariates - age, education, age at start of any music lessons, musical training, and musical listening - are added to a simple model that uses only gender. This analysis was run on the main study data. *R*^2^ values lower in full model due to missing observations (some participants opted to not report information for extra covariates).

| Task | $\eta^2$ of<br>gender in<br>simple<br>model | $R^2$ simple<br>model | partial $\eta^2$<br>of gender<br>in full<br>model | $R^2$ full<br>model |
| --- | --- | --- | --- | --- |
| Mistuning Perception | 0.001 | 0.076 | 0.002 | 0.002 |
| Melodic Discrimination | 0.001 | 0.078 | 0.000 | 0.000 |
| Beat Alignment | 0.004 | 0.038 | 0.003 | 0.003 |
| General Music Perception Ability | 0.000 | 0.119 | 0.000 | 0.000 |

**SI Table 3.**
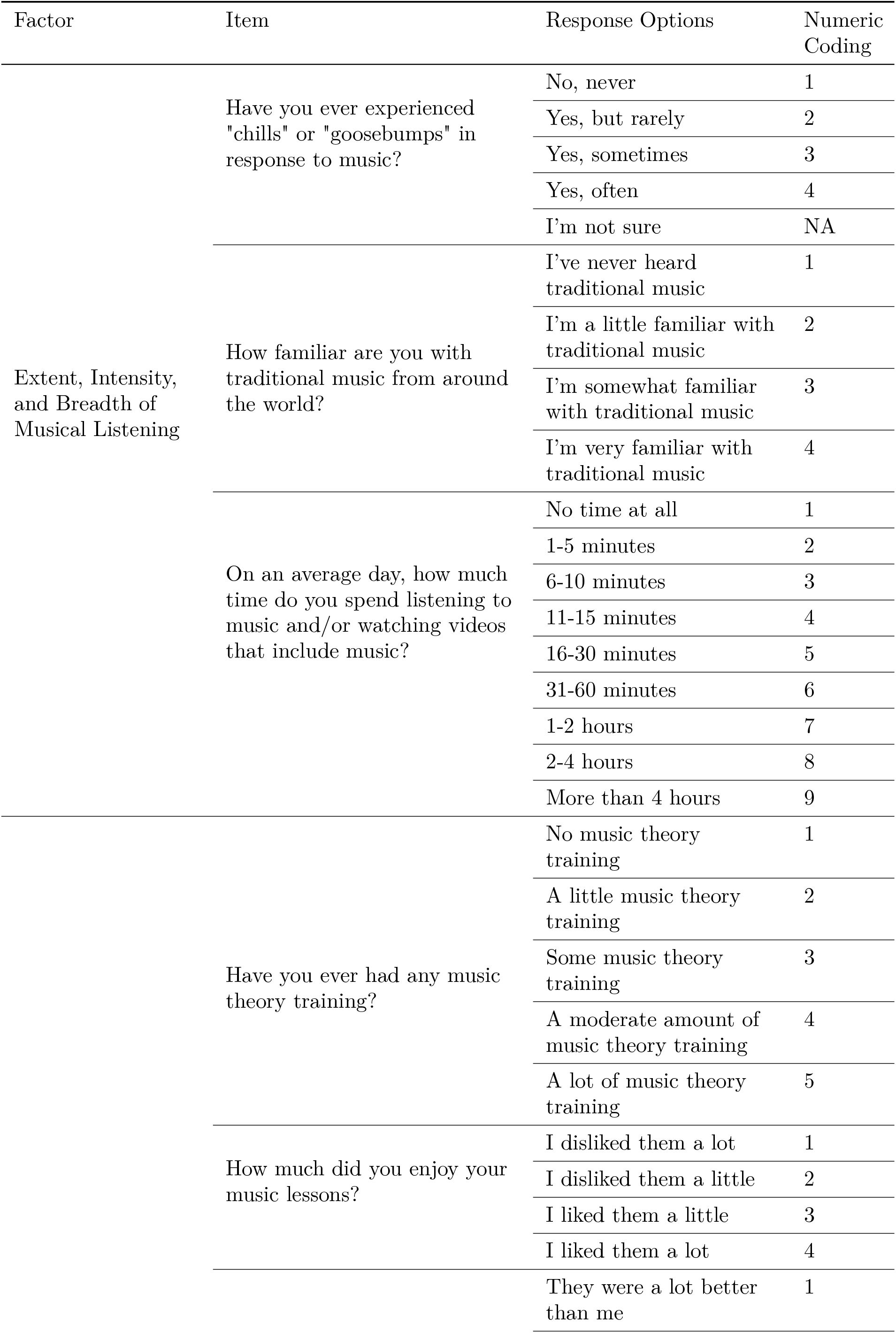
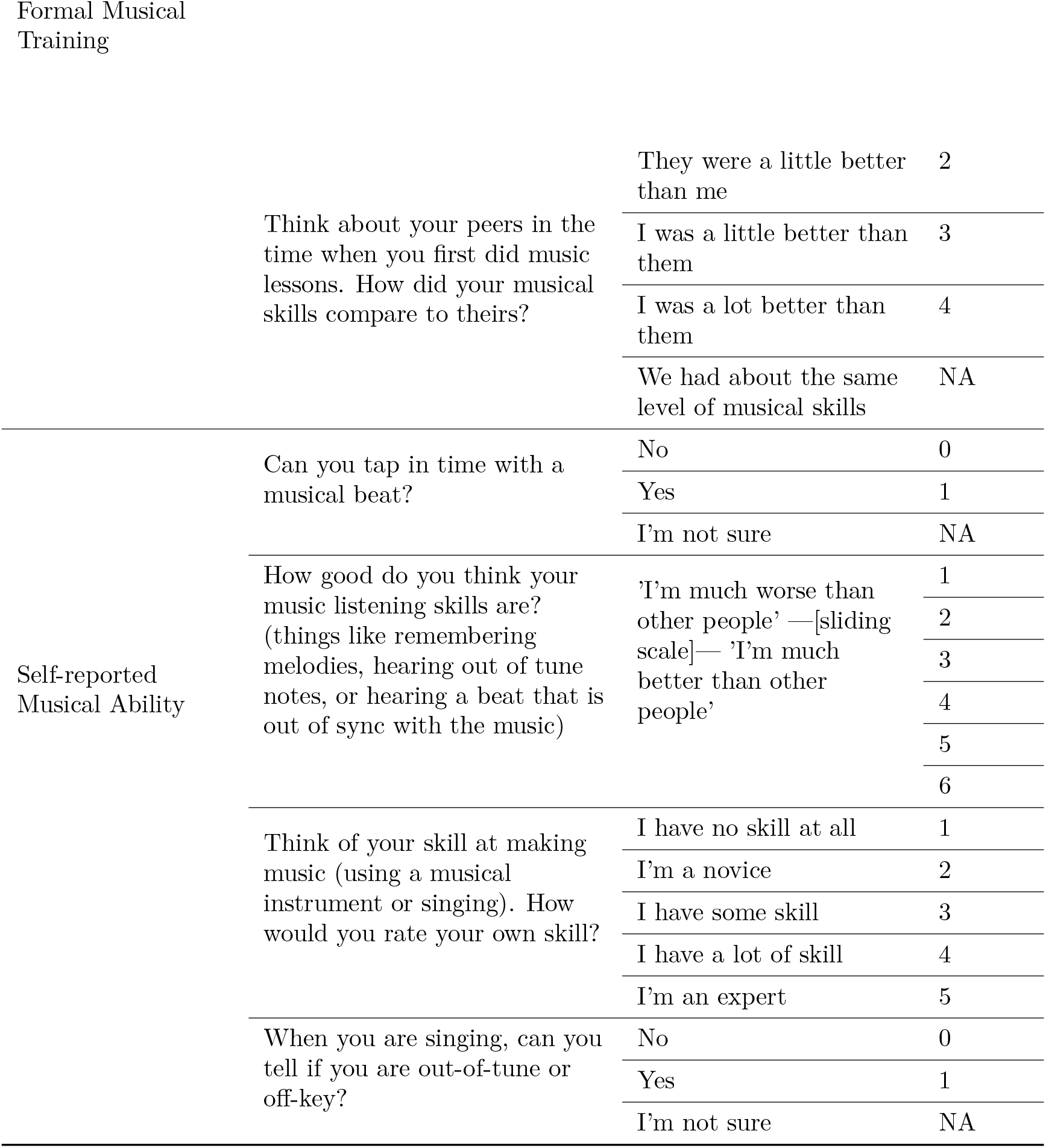
Items. Details of item verbal prompts, response options, and numeric coding of responses.

**SI Table 4.**
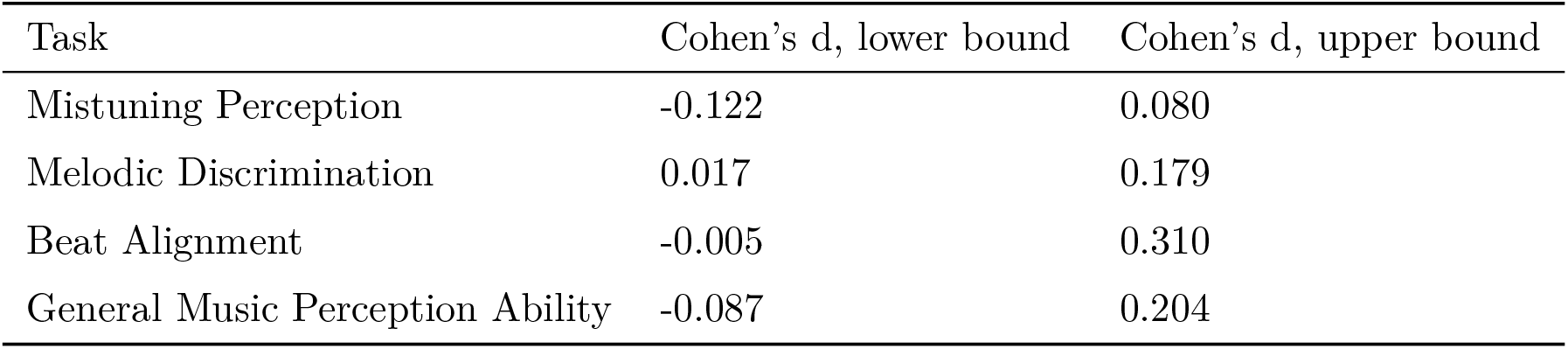
Range of Cohen’s d observed in tree models. Tree models estimated the magnitude of sex differences in music perception tests in different subgroups of participants in the main study, as partitioned on participant age, education, start age of music lessons, and factors musical training and musical listening. Negative scores indicate a male advantage.

## Supplementary Figures

**SI Figure S1.**
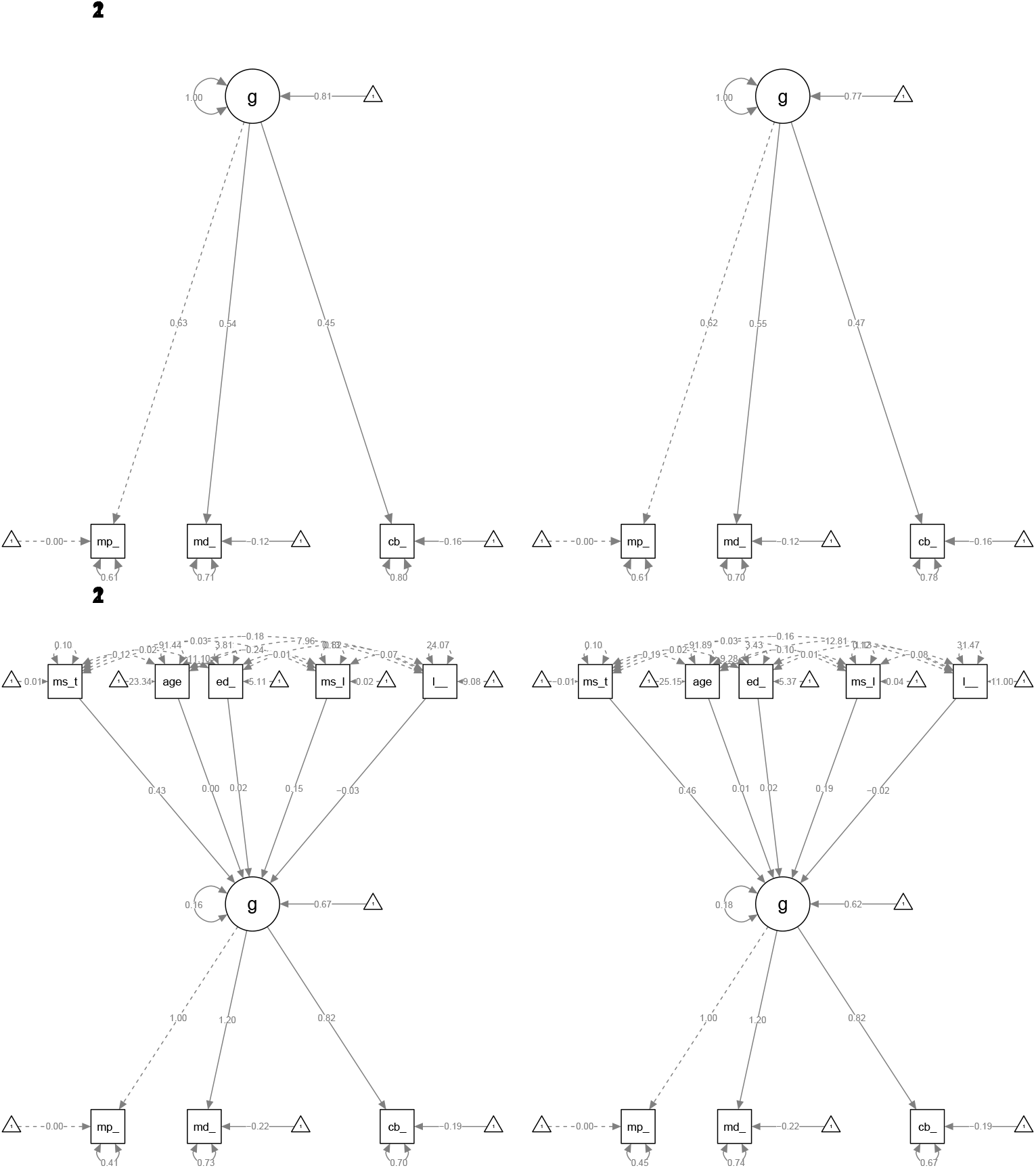
Structural Equation Models. Top row shows gender-only models, bottom row shows all-covariate models. Left panels in each row show the models for female participants; right panels show these models for males.

**SI Figure S2.**
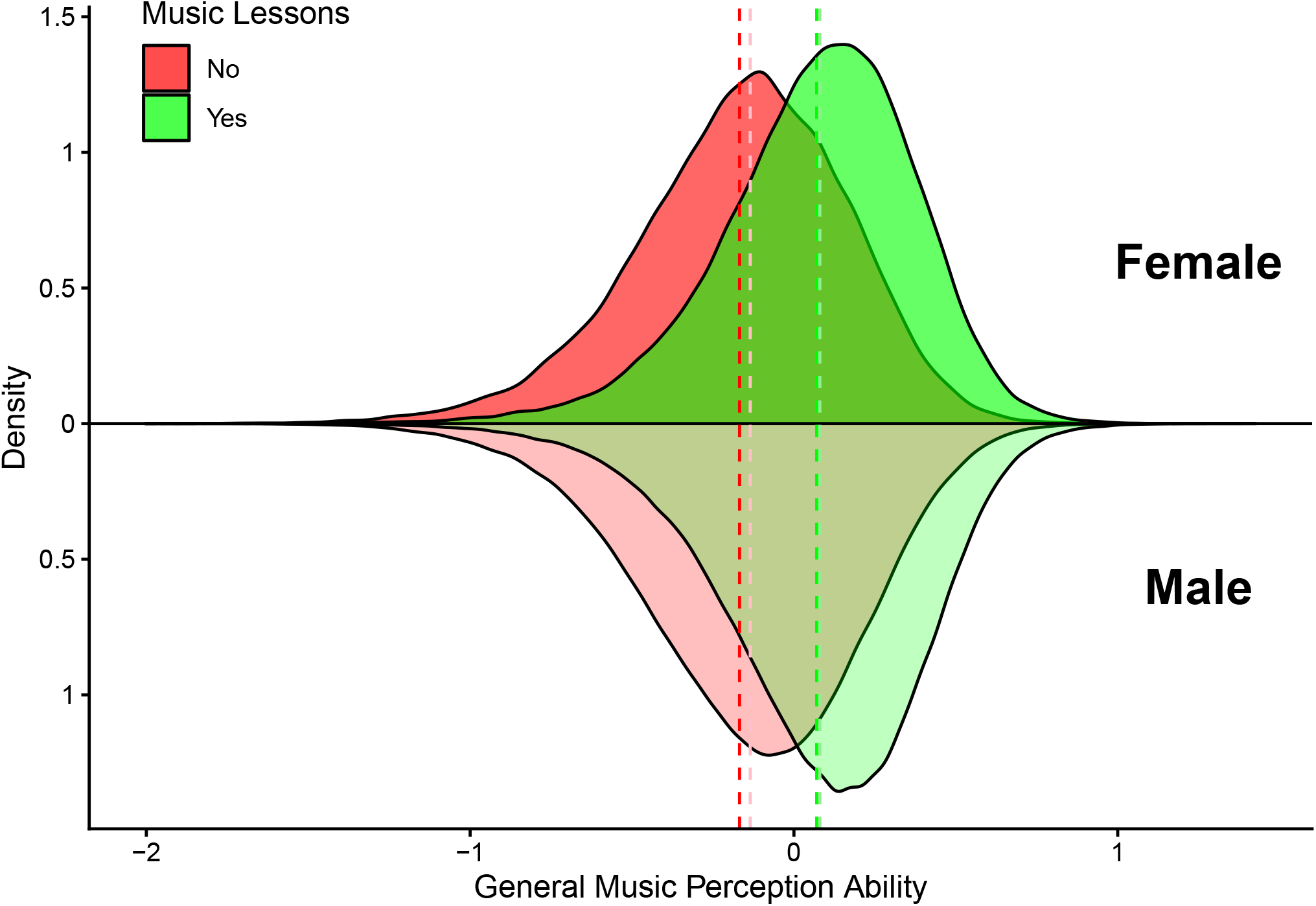
Participants who had taken music lessons had higher musical ability than paticipants who had not. There was a difference in general music perception ability scores of d=0.684 between the those who self report having had or not had music lessons in their life. This same pattern manifested similarly among men and women. Among those who reported no music lessons, we observed a sex difference of Cohen’s d = -0.099. Bolder colours for distributions and lines indicate female distributions; dashed lines indicate mean general music perception ability, for each subgroup of gender*music lessons.

## Notes

https://osf.io/3v27y/

### Competing Interest Statement

The authors have declared no competing interest.

### Summary of Updates

Three replication studies. The main finding was replicated in two follow-up studies where participant recruitment was outsourced to a third party that built balanced (gender, age, socioeconomic status) participant pools in the US and UK; and replicated in a third replication study that was a direct replication of the original finding.

